# Phenotypic Characterization of indigenous goat population in Southern, Ethiopia

**DOI:** 10.1101/2022.02.16.480639

**Authors:** Teshager Muluneh, Wondimagegn Tadesse

## Abstract

A study was conducted at Abaya and Yirgachafe districts to characterize indigenous goat types phenotypically. Data were collected through field measurements and visual observation of qualitative traits. Totally 540 goats were used for metric and morphometric measurement. Results of the study revealed that the goat populations found in Abaya and Yirgachafe district were different characteristics which are physically Abaya goats were closest with Arsi-Bale whereas yirgachafee with Woyto-Guji which are mostly distributed goat breeds in southern Ethiopia. The dominant coat color pattern in study area was plain, patchy, and spotted with proportions of 55.19, 37.04, and 7.78% and 46.67, 38.89, and 14.44% in Abaya and Yirgachafee district respectively. A strong and positive correlation(r = 0.83, 0.76) was observed between heart girth and body weight for male and female goat populations respectively. Generally, the indigenous goat population has its own difference in its morphological and morphometric traits. Traits have their own economic contribution. Therefore, identifying these important traits for further genetic improvements, conservation and sustainable utilization of the genetic resources of the diversified goat population is important.

## Introduction

Goats (Capra hircus) contribute significantly to the livelihood of resource-poor farmers in Ethiopia. Goats have a short reproductive cycle hence high multiplication rate as compared to large ruminants, which is ideal for poverty alleviation providing income, meat, milk, skin and manure, as aliving bank against the various environmental hazards (crop failure, drought and flooding) and have serve for socio-cultural values for diverse traditional communities (Tesfahun et al., 2017, Hirpa and Abebe, 2008)

Ethiopian goats are classified in to eight genetically diverse breeds which adapted to a range of environments from arid lowlands (the pastoral and agro-pastoral production system) to the humid highlands (mixed farming systems) (Tucho and Tesfaye, 2004). Ethiopia has about 32.74 million goats, of which about 70.49 percent are females and 29.51 percent are males and with respect to breed, almost all of the goats are indigenous breeds, which account about 99.97 % (CSA, 2018).

Iindigenous goat populations generally dominate the goat flocks in Ethiopia and have developed certain valuable genetic traits such as ability to perform better under low input condition and climatic stress, tolerance to infectious diseases and parasites as well as heat stresses (Hassen, 2012). Their morphological differences have important socio-cultural and economic values to the Ethiopian communities; as a result, most farmers have specific consideration and choices for goat coat colors followed by body sizes. For instance, black coat colored goat is less preferred in the Amhara Region and beyond (Hassen, 2012).

A systematic description/characterization of the goat types and management systems should be considered as prerequisite for planning the rational use of indigenous goat resources. In addition breed characterization is the first step in the urgent task of genetic resource management and conservation of goat on the risk status (FAO, 2012). Breed characterization can be done through performance evaluation, phenotypic characterization and DNA molecular characterization (FAO, 2015) which provide comprehensive database information of variation among the goat populations as to which of the populations represent homogenous populations and which of them are genetically distinct, these all information would generate understanding of the goat type.

Based on this genetic characterization of Arsi-Bale and Woyito-Guji breeds which is distribution overlap in the current study area of Abaya and Yirgachefe districts. Therefore, even though genetic characterization for Arsi-Bale and Woyito-Guji breeds have been done, which are distributed in southern part of Ethiopia, due to overlapping of the distribution of these two breeds in the study area the present phenotypic characterization of indigenous goat was initiated. Despite the studies done, information on phenotypic characteristics and production systems of some indigenous goat populations is still scanty. Besides, there was little intervention works so far on the improvement of production and productivity of local goat breeds in the area. Also farmers practice traditional type of goat production system in the area.

## Material and Methods

### Description of the Study Areas

The study was conducted in two National Regional States of Ethiopia: Oromiya regional states (Abaya district) in West Guji zone and South nation nationalty and peoples region (Yirgachafee district) of Gedio zone.Eventhogh only adiminstrative demarkation makes, at different region unless districts are interborder and located southern part of Ethiopia. Abaya is one of the districts in the Oromia Region of Ethiopia. It is part of former Gelana-Abaya district that was divided later on as Abaya and Gelana districts. This district is located between latitude of **5**^0^ 45**’0’’**N-6^**0**^ 45**’00’’**N and longitude of 37^*0*^44’ 00”E-38^**0**^20**’** 00’’E. It is part of the West Guji zone, Abaya was bordered on the south by Bule Hora and on the west, north and east by Southern nations, nationalities, and peoples region (SNNP) and Lake Abaya, on the western. (District Agriculture and Natural resource office, 2019).

Yirgachafee is also one of district of Gedeo zone, in SNNP region of Ethiopia. This study area is located at about 395 km south of Addis Ababa, capital city of Ethiopia and 124km from Hawassa. Yirgachafee is bordered on the south by Kochore districts, west by the Abaya district of West Guji zone, and north by Wenago, east by Bule and southeast by Gedeb. The district is located at latitude between 6^0^4’00’’N-6^0^15 00’’N and longitude of 38^0^10’00’’E-38^0^ 20’00’’E. (District Agriculture and Natural resource office, 2019).

### Sampling techniques, sample size and data collection

Purposive sampling techniques were applied to select both study districts and Kebeles based up on the size of goat population obtained from respective agriculture and natural resource office. Each household from kebele was selected randomly from listed households based on year and experience of goat rearing at least two year. The site selection and the household baseline surveys were conducted from 1 September to begning of December 2019.

A total of 540 (162 males and 378 female) goat were sampled for quantitative (Body weight (BW) and linear body measurments (LBM) like height at wither (HW), body length (BL), heart girth (HG), ear length (EL), pelvic width (PW), chest depth (CD) and scrotal circumference (SC) for male using measuring tape in level ground and weight of goat was taken using 50 kg spring balance using sack bag by hanging and ground balance for those who owen coffee ground beambalence. Linear body measurements were taken on goats which have one and above pair of permanent (1PPI, 2PPI, 3PPI and **>**4PPI) and qualitative (coat color pattern, coat color type, head profile, back profile, rump profile, ear orientation, horn (presence, absence, shape and orientations), hair type, toggle, ruff; beard) using visual observations based on breed description list of FAO (2012).

### Statistical Data Analysis

All data gathered during the study period were coded and recorded in Microsoft Excel 97-2003.The data were analyzed by SAS version 9.2 (2008). General linear model procedure (PROC GLM) of SAS was used for both metric and morphometric trait analysis. Tukey’s comparison test was used to compare the sub factor brought significant difference. Descriptive statistics were also used to describe the results as percentages for all districts.

Body weight and LBMs for both sexes was analysed using follwing mode

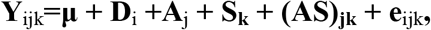

**where;**

**Y**_ijk_ = the observed value of trait of interest,

μ = overall mean

D_***i***_ = the effect of i^th^ district (i=1, 2)

A_j_ = the effect of the j^th^ age (dentition class) (j=1, 2, 3, 4 pairs of permanent incisor)

S_k_ = the effect of k^th^ sex (k = male, female),

AS_jk_ = Interaction effect of j^th^ age (dentition class) and k^th^ sex,

e_ijk_ = the residual random error.

The model employed for analyses of scrotal circumference and length:

**Y**_ijk_**=μ + D**_*i*_ **+ A**_j_ + **e**_ijk_, **Where;**

**Y**_ijk_ = the observed value of scrotal circumference and length,

μ = overall mean,

D_***i***_ = the effect of i^th^ district (i=1,2)

A_j_ = the effect of the j^th^ age (dentition class) (j=1,2,3,4),

e_ijk_ = the residual random error

## Result

### Qualitative traits

In this particualr study, the overall coat color patterns for both sexes were plain (46.67% and patchy/pied (38.89). The overall observed coat color type for both sex were predominantly red (16.66%), followed by white dominant and grey (13.70%) and white (12.22%) in Abaya, whereas black dominant (14.81%), fawn (13.70%) and white dominant (13.33%) in Yirgachafee. Goat population in study district was unified with smooth hair followed by glossy and curly rough hair type with 40.74, 24.07 and 20 % in Abaya where as 38.89, 23.70 and 20.7% smooth, long straight and curly rough in Yirgachafee respectively. In both districts 90-94% of the goat population had horn.

**Table 1.**
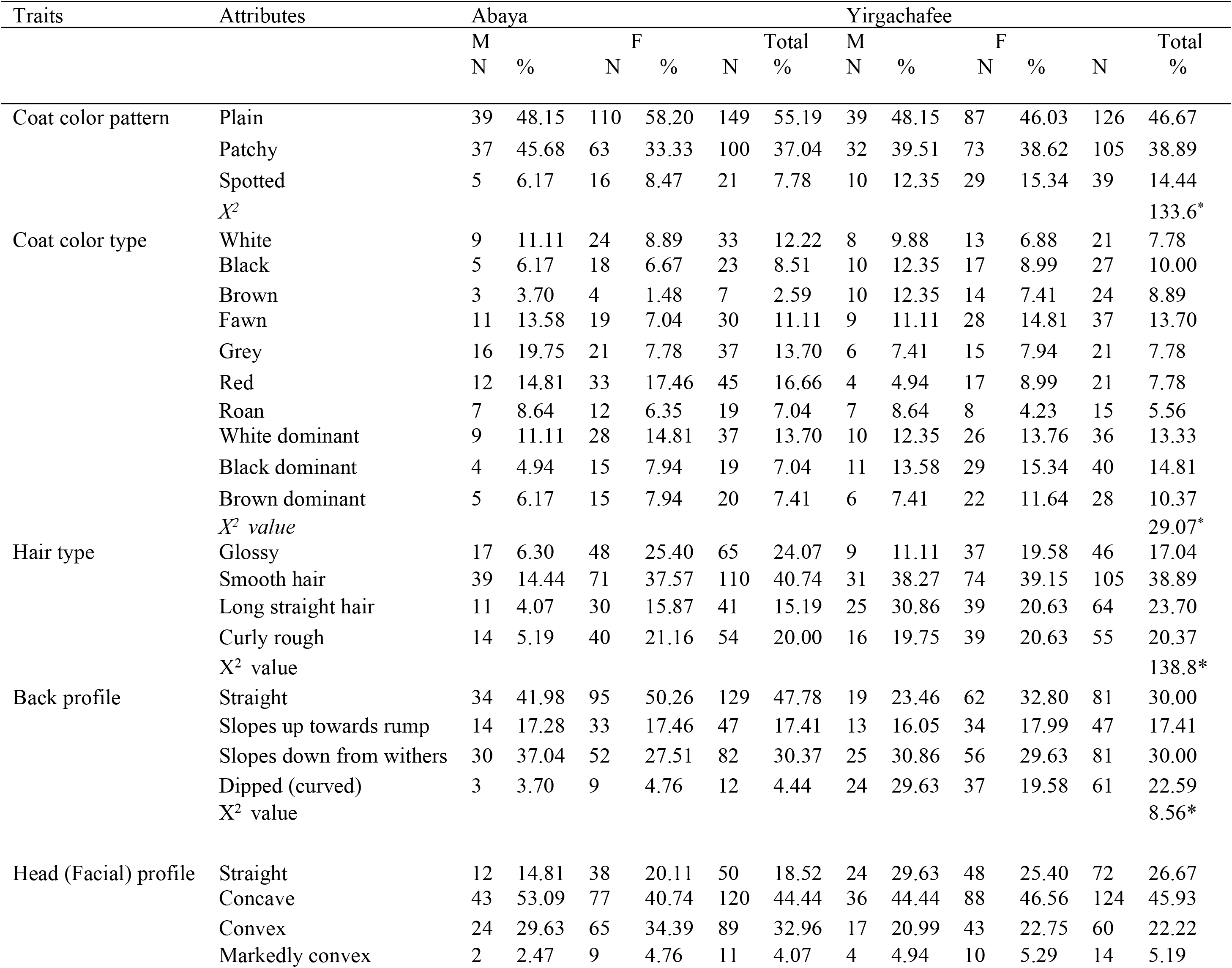
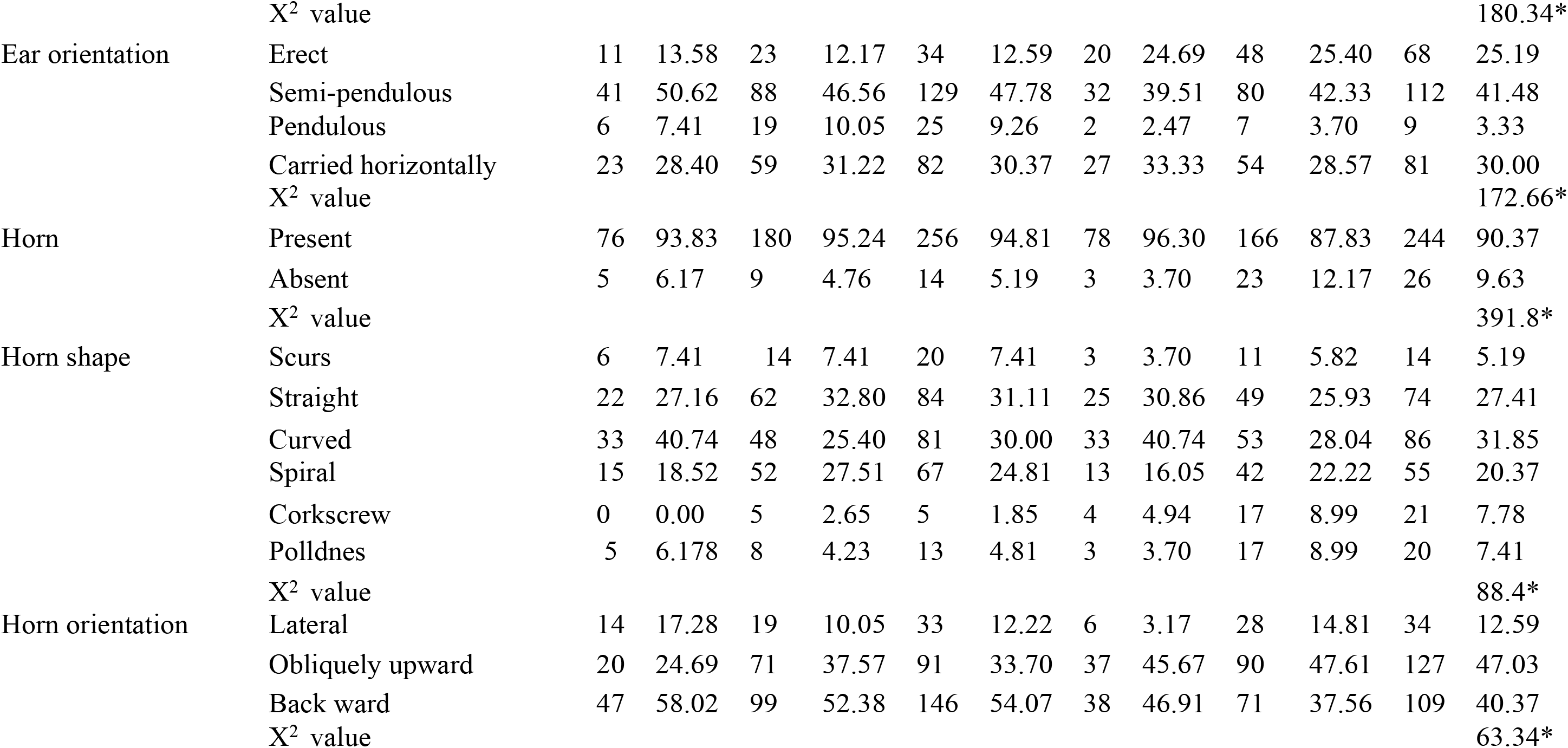
Description on qualitative traits of goat

| Traits | Attributes | Abaya |  |  |  |  |  | Yirgachafee |  |  |  |  |  |
| --- | --- | --- | --- | --- | --- | --- | --- | --- | --- | --- | --- | --- | --- |
|  |  | M |  | F |  |  | Total | M |  | F |  |  | Total |
|  |  | N | % | N | % | N | % | N | % | N | % | N | % |
| Coat color pattern | Plain | 39 | 48.15 | 110 | 58.20 | 149 | 55.19 | 39 | 48.15 | 87 | 46.03 | 126 | 46.67 |
|  | Patchy | 37 | 45.68 | 63 | 33.33 | 100 | 37.04 | 32 | 39.51 | 73 | 38.62 | 105 | 38.89 |
|  | Spotted | 5 | 6.17 | 16 | 8.47 | 21 | 7.78 | 10 | 12.35 | 29 | 15.34 | 39 | 14.44 |
| | $\chi^2$ | | | | | | | | | | | | 133.6* |
| Coat color type | White | 9 | 11.11 | 24 | 8.89 | 33 | 12.22 | 8 | 9.88 | 13 | 6.88 | 21 | 7.78 |
|  | Black | 5 | 6.17 | 18 | 6.67 | 23 | 8.51 | 10 | 12.35 | 17 | 8.99 | 27 | 10.00 |
|  | Brown | 3 | 3.70 | 4 | 1.48 | 7 | 2.59 | 10 | 12.35 | 14 | 7.41 | 24 | 8.89 |
|  | Fawn | 11 | 13.58 | 19 | 7.04 | 30 | 11.11 | 9 | 11.11 | 28 | 14.81 | 37 | 13.70 |
|  | Grey | 16 | 19.75 | 21 | 7.78 | 37 | 13.70 | 6 | 7.41 | 15 | 7.94 | 21 | 7.78 |
|  | Red | 12 | 14.81 | 33 | 17.46 | 45 | 16.66 | 4 | 4.94 | 17 | 8.99 | 21 | 7.78 |
|  | Roan | 7 | 8.64 | 12 | 6.35 | 19 | 7.04 | 7 | 8.64 | 8 | 4.23 | 15 | 5.56 |
|  | White dominant | 9 | 11.11 | 28 | 14.81 | 37 | 13.70 | 10 | 12.35 | 26 | 13.76 | 36 | 13.33 |
|  | Black dominant | 4 | 4.94 | 15 | 7.94 | 19 | 7.04 | 11 | 13.58 | 29 | 15.34 | 40 | 14.81 |
|  | Brown dominant | 5 | 6.17 | 15 | 7.94 | 20 | 7.41 | 6 | 7.41 | 22 | 11.64 | 28 | 10.37 |
| | $\chi^2$ value | | | | | | | | | | | | 29.07* |
| Hair type | Glossy | 17 | 6.30 | 48 | 25.40 | 65 | 24.07 | 9 | 11.11 | 37 | 19.58 | 46 | 17.04 |
|  | Smooth hair | 39 | 14.44 | 71 | 37.57 | 110 | 40.74 | 31 | 38.27 | 74 | 39.15 | 105 | 38.89 |
|  | Long straight hair | 11 | 4.07 | 30 | 15.87 | 41 | 15.19 | 25 | 30.86 | 39 | 20.63 | 64 | 23.70 |
|  | Curly rough | 14 | 5.19 | 40 | 21.16 | 54 | 20.00 | 16 | 19.75 | 39 | 20.63 | 55 | 20.37 |
| | $\chi^2$ value | | | | | | | | | | | | 138.8* |
| Back profile | Straight | 34 | 41.98 | 95 | 50.26 | 129 | 47.78 | 19 | 23.46 | 62 | 32.80 | 81 | 30.00 |
|  | Slopes up towards rump | 14 | 17.28 | 33 | 17.46 | 47 | 17.41 | 13 | 16.05 | 34 | 17.99 | 47 | 17.41 |
|  | Slopes down from withers | 30 | 37.04 | 52 | 27.51 | 82 | 30.37 | 25 | 30.86 | 56 | 29.63 | 81 | 30.00 |
|  | Dipped (curved) | 3 | 3.70 | 9 | 4.76 | 12 | 4.44 | 24 | 29.63 | 37 | 19.58 | 61 | 22.59 |
| | $\chi^2$ value | | | | | | | | | | | | 8.56* |
| Head (Facial) profile | Straight | 12 | 14.81 | 38 | 20.11 | 50 | 18.52 | 24 | 29.63 | 48 | 25.40 | 72 | 26.67 |
|  | Concave | 43 | 53.09 | 77 | 40.74 | 120 | 44.44 | 36 | 44.44 | 88 | 46.56 | 124 | 45.93 |
|  | Convex | 24 | 29.63 | 65 | 34.39 | 89 | 32.96 | 17 | 20.99 | 43 | 22.75 | 60 | 22.22 |
|  | Markedly convex | 2 | 2.47 | 9 | 4.76 | 11 | 4.07 | 4 | 4.94 | 10 | 5.29 | 14 | 5.19 |

### Live Body Weight and Linear Measurement

In the study area, overall mean of live body weight, heart girth, height at wither, body length, scrotal length, scrotal circumference were 31.1 kg, 71.8cm, 58.9cm, 52.2cm, 22.87cm, 16.10 cm, respectively.

**Location effect:-** There was significant difference (p<0.05) in body weight and all linear body measurements except HL, FC and FH between both districts.

**Sex effect**:- Heart girth, bodyweight, pelvic width, chestdepth and neck length was significantly affected by sex.

**Age effect**:- Age has significant (p<0.05) differences for all linear body measurements except forecanon circumstance and fore canon height. In this study body weight (BW) and some linear body measurements significant difference were observed among age groups. The scrotal circumference and length was also significantly (p<0.05) affected by age.

**Sex by Age group:-** The interaction of sex and age group was not significantly (p>0.05) different for body weight and other body measurements except body length, chest depth and fore canon circumstance. Bucks at age category of 1PP and 4PP are higher in body length than does in the respective age.

**Table 2.**
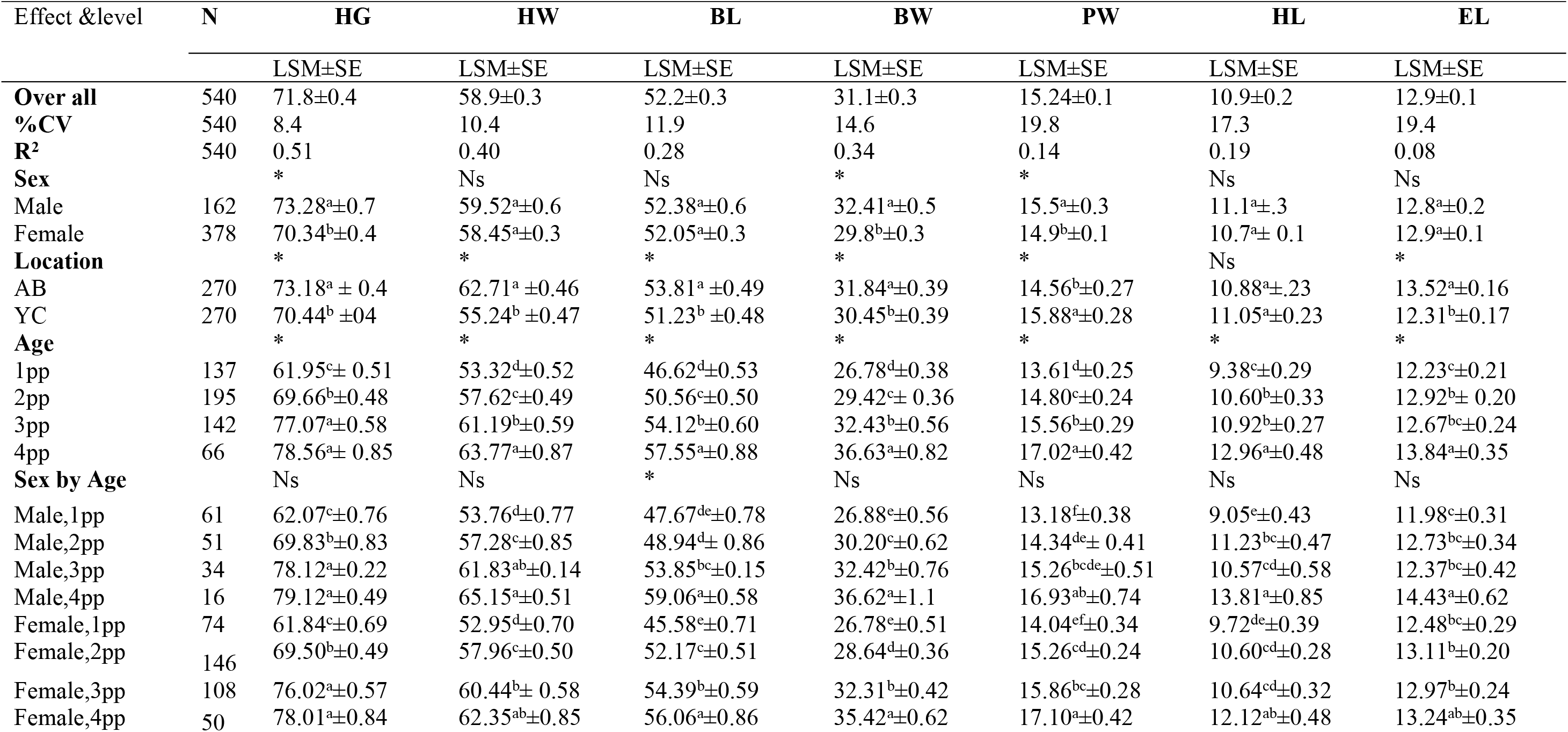
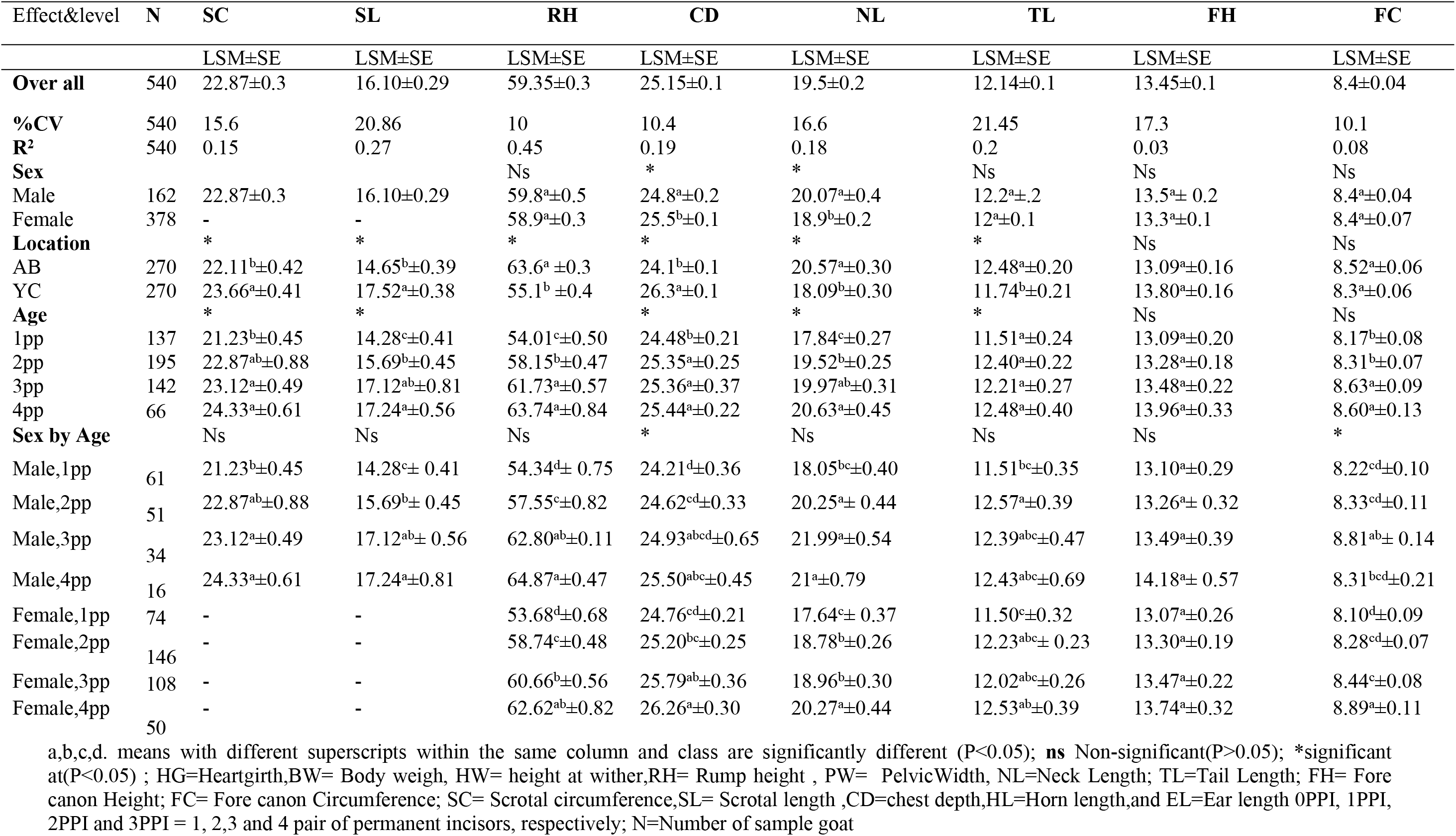
Live body weight and linear body measurement of goat.

### Stepwise Discriminate Analysis

Stepwise discriminate analysis procedure identified five variables for buck and these are rump height (RH), chest depth (CD), forcanon height (FH), height at wither (HW) and Scrotal length (SL) and six for doe rump height (RH), chest depth (CD), forcanone height (FH), body weight (BW), neck length (NL) and ear length (EL) as most significant discriminating traits.

**Table 3.**
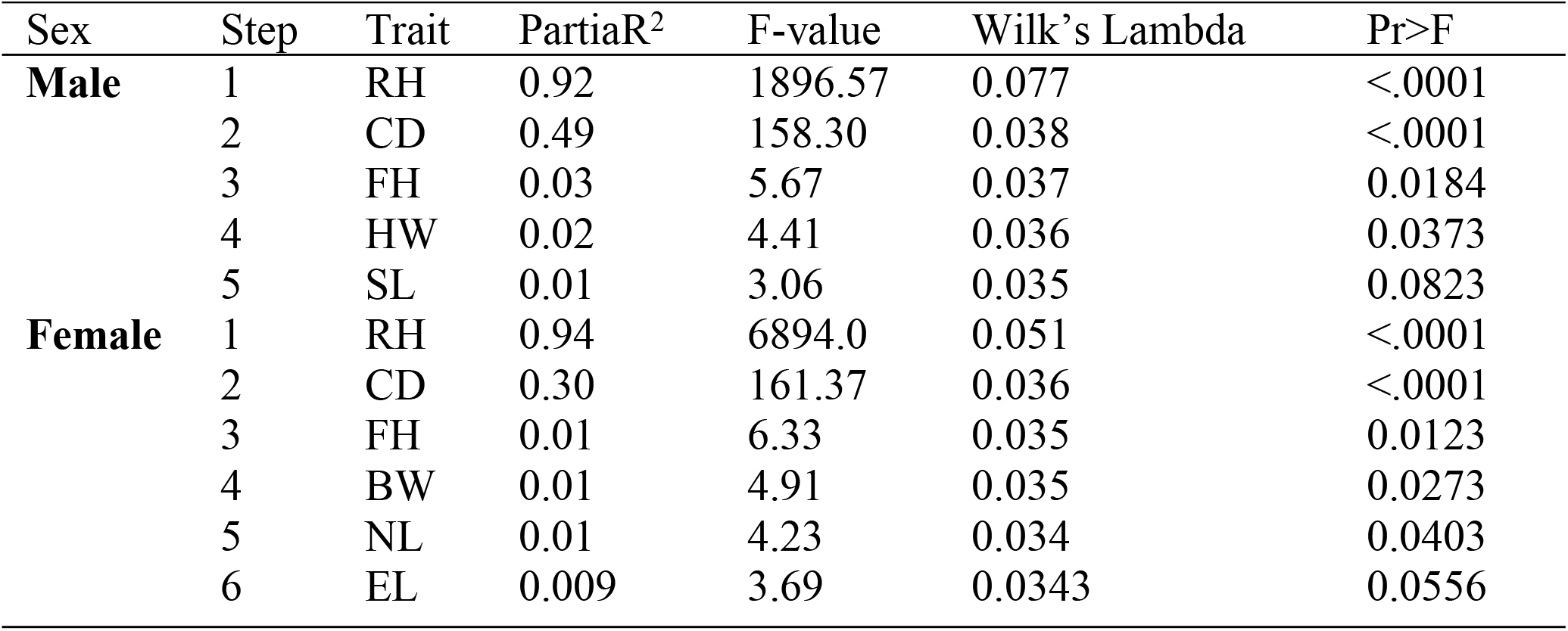
Summary of stepwise selection of traits for buck and does

| Sex | Step | Trait | PartiaR<sup>2</sup> | F-value | Wilk's Lambda | Pr>F |
| --- | --- | --- | --- | --- | --- | --- |
| **Male** | 1 | RH | 0.92 | 1896.57 | 0.077 | <.0001 |
|  | 2 | CD | 0.49 | 158.30 | 0.038 | <.0001 |
|  | 3 | FH | 0.03 | 5.67 | 0.037 | 0.0184 |
|  | 4 | HW | 0.02 | 4.41 | 0.036 | 0.0373 |
|  | 5 | SL | 0.01 | 3.06 | 0.035 | 0.0823 |
| **Female** | 1 | RH | 0.94 | 6894.0 | 0.051 | <.0001 |
|  | 2 | CD | 0.30 | 161.37 | 0.036 | <.0001 |
|  | 3 | FH | 0.01 | 6.33 | 0.035 | 0.0123 |
|  | 4 | BW | 0.01 | 4.91 | 0.035 | 0.0273 |
|  | 5 | NL | 0.01 | 4.23 | 0.034 | 0.0403 |
|  | 6 | EL | 0.009 | 3.69 | 0.0343 | 0.0556 |

### Prediction of Body Weight from Linear Body Measurements

All body measurements were fitted into the model and through elimination procedures, the optimum model was identified heart girth (HG), height at wither (HW), body length (BL), pelvic width (PW) and horn length (HL) for male where as body length (BL), heart girth (HG), height at wither (HW) and pelvic width (PW) was the best fitted model for female. Heart girth, hight at wither, body length, pelvic width and horn length were include in the model in order of importance and they account 63% of the total variability and heart girth alone accounts for 39% variation in the body weight for buck.

In female sampled goat population four variables were positively contributing to the prediction of model which include heart girth, hight at wither, body length and pelvic width were fitted as first, second, third and fourth which account 84% of total variability and heart girth alone also acounts 51% of variation in body weight. The predicted equation of body weight for both male and female are presented below:**-** Body Weight = -5.98 +0.17 HG + 0.25 HW + 0.24 BL + 0.06 PW +0.05 HL for Male Body Weight = 12.25 + 0.15 HG +0.01HW+ 0.16 BL + 0.15 PW for Female where HG, HW, BL, PW and HL explanatory or independent variables.

**Table 7.**
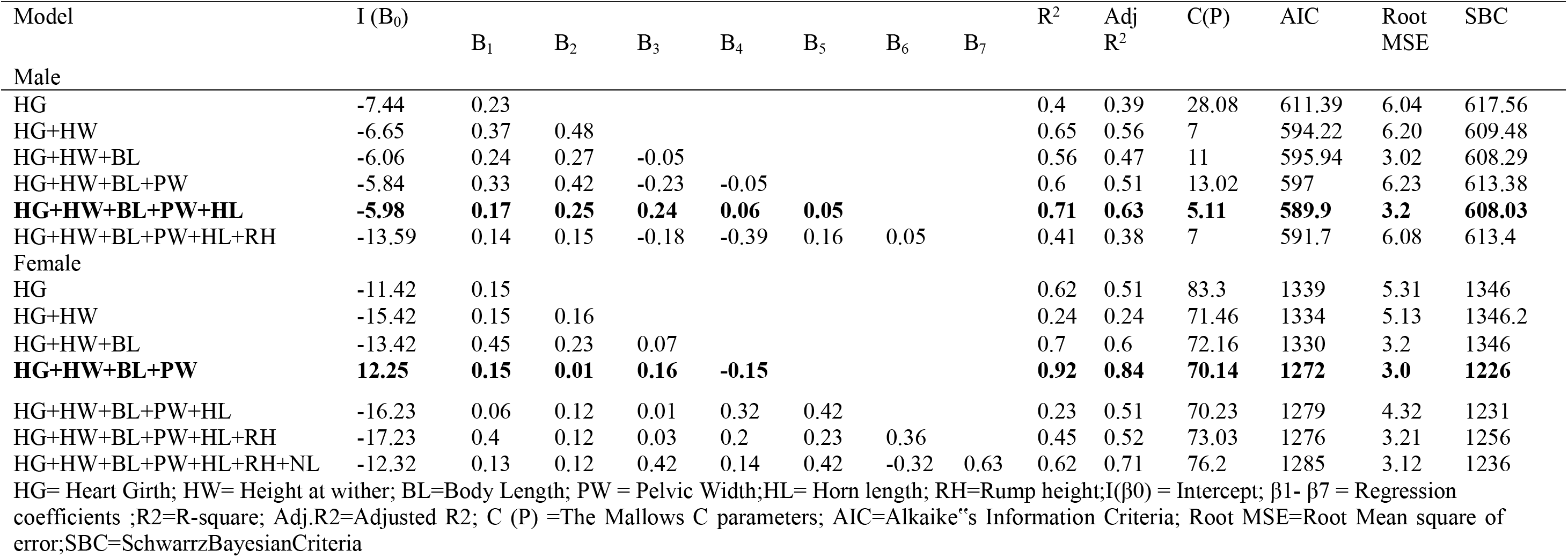
Multiple linear regression analysis of live body weight on different LBMs for male and female goat in all age groups

| Model | I (B<sub>0</sub>) | B<sub>1</sub> | B<sub>2</sub> | B<sub>3</sub> | B<sub>4</sub> | B<sub>5</sub> | B<sub>6</sub> | B<sub>7</sub> | R<sup>2</sup> | Adj R<sup>2</sup> | C(P) | AIC | Root MSE | SBC |
| --- | --- | --- | --- | --- | --- | --- | --- | --- | --- | --- | --- | --- | --- | --- |
| Male |  |  |  |  |  |  |  |  |  |  |  |  |  |  |
| HG | -7.44 | 0.23 |  |  |  |  |  |  | 0.4 | 0.39 | 28.08 | 611.39 | 6.04 | 617.56 |
| HG+HW | -6.65 | 0.37 | 0.48 |  |  |  |  |  | 0.65 | 0.56 | 7 | 594.22 | 6.20 | 609.48 |
| HG+HW+BL | -6.06 | 0.24 | 0.27 | -0.05 |  |  |  |  | 0.56 | 0.47 | 11 | 595.94 | 3.02 | 608.29 |
| HG+HW+BL+PW | -5.84 | 0.33 | 0.42 | -0.23 | -0.05 |  |  |  | 0.6 | 0.51 | 13.02 | 597 | 6.23 | 613.38 |
| **HG+HW+BL+PW+HL** | **-5.98** | **0.17** | **0.25** | **0.24** | **0.06** | **0.05** |  |  | **0.71** | **0.63** | **5.11** | **589.9** | **3.2** | **608.03** |
| HG+HW+BL+PW+HL+RH | -13.59 | 0.14 | 0.15 | -0.18 | -0.39 | 0.16 | 0.05 |  | 0.41 | 0.38 | 7 | 591.7 | 6.08 | 613.4 |
| Female |  |  |  |  |  |  |  |  |  |  |  |  |  |  |
| HG | -11.42 | 0.15 |  |  |  |  |  |  | 0.62 | 0.51 | 83.3 | 1339 | 5.31 | 1346 |
| HG+HW | -15.42 | 0.15 | 0.16 |  |  |  |  |  | 0.24 | 0.24 | 71.46 | 1334 | 5.13 | 1346.2 |
| HG+HW+BL | -13.42 | 0.45 | 0.23 | 0.07 |  |  |  |  | 0.7 | 0.6 | 72.16 | 1330 | 3.2 | 1346 |
| **HG+HW+BL+PW** | **12.25** | **0.15** | **0.01** | **0.16** | **-0.15** |  |  |  | **0.92** | **0.84** | **70.14** | **1272** | **3.0** | **1226** |
| HG+HW+BL+PW+HL | -16.23 | 0.06 | 0.12 | 0.01 | 0.32 | 0.42 |  |  | 0.23 | 0.51 | 70.23 | 1279 | 4.32 | 1231 |
| HG+HW+BL+PW+HL+RH | -17.23 | 0.4 | 0.12 | 0.03 | 0.2 | 0.23 | 0.36 |  | 0.45 | 0.52 | 73.03 | 1276 | 3.21 | 1256 |
| HG+HW+BL+PW+HL+RH+NL | -12.32 | 0.13 | 0.12 | 0.42 | 0.14 | 0.42 | -0.32 | 0.63 | 0.62 | 0.71 | 76.2 | 1285 | 3.12 | 1236 |

HG= Heart Girth; HW= Height at wither; BL=Body Length; PW = Pelvic Width;HL= Horn length; RH=Rump height;I(β<sub>0</sub>) = Intercept; β<sub>1</sub>- β<sub>7</sub> = Regression coefficients ;R<sup>2</sup>=R-square; Adj.R<sup>2</sup>=Adjusted R<sup>2</sup>; C (P) =The Mallows C parameters; AIC=Alkaike's Information Criteria; Root MSE=Root Mean square of error;SBC=SchwarzBayesianCriteria

## Discussion

In this particualr study, the overall coat color patterns for both sexes were plain (46.67%) and patchy/pied (38.89%) in both districts. Different authors reported different coat colar patterns. For instances, Hassen (2012) reported spotted and patchy for North Amhara goat population while it was plain coat color patterns for Arsi-Bale goats (Belete, 2013). Zergaw and Dessie (2014) also reported similar coat color patterns for Woyto Guji goats in SNNP indicating that Abaya and Yirgachafee goats share common coat color patterns with Woyto Guji and Arsi-Bale goats probably as a result of gene flow between these two neighboring populations.

Information on body and testicle size of specific goat breed at constant age has paramount importance in the selection of genetically superior animals for production and reproduction purpose. The fact that physical linear traits have medium-to high heritability and are well correlated with BW indicates their importance for effective selection (Magnabosco *et al*.,2002).

**Location effect:-** There was significant difference (p<0.05) in body weight and all linear body measurements except HL, FC and FH between both districts. The reason for the significant difference of live body weghit and other linear body measurment accros districts was due to availabilty of feed in the free browsing areas of Abaya district. The current finding of body weight of sampled goat was comparable with report of Gatew (2015) who reported for Bati and Borena goat population but higher than short ear somali goat 33.97±0.49kg, 31.49±0.23kg and 24.67±028kg respectivly.

**Sex effect**:- Heart girth, bodyweight, pelvic width, chestdepth and neck length was significantly affected by sex. In species having sexual dimorphism, the two sex may vary in color, size, or some other traits (Isaac, 2005). The same was true in this study where males were superior than females in body weight, heart girth, pelvic width, chest depth and neck length. The sex related differences might be partly a function of the sex differential hormonal effect on growth (Semakula **et al**., 2010).

The effect of age shows that as age increase body weight and other linear body measurements were increases. According to Yoseph (2007) reported that body size and shape of animal rises until the animal reaches the optimal growth. Maximum value was observed in age class of three and four as compared to one and two. The present finding agree with that of Takele (2016) who report body weight and linear body measurement were increased as age of animal became old. The scrotal circumference and length was also significantly (p<0.05) affected by age. The size of scrotal circumference and length increases as age increase from one pair of permanent incisor to fourth pair of permanent incisors. This finding is consistent with the study Yoseph (2007) described that breed, age and their interaction significantly affected by BW, body condition score (BCS), scrotal circumference (SC) and testicular weight (TW).

Scrotal circumference is the most heritable components of fertility that should be included for evaluation of breeding soundness (Yoseph, 2007). The scrotal circumference at the age of 3PP in this study was lower than Bati and Borena (27cm) bucks but comparable with Short eared Somali bucks (25cm) (Gatew, 2015). The observed difference of SC between bucks in the study area and Bati and Borena bucks were may be due to the higher body weight exhibited in Bati and Borena Bucks. Besides, the study Yoseph (2007) explained that scrotal traits were directly influenced by agro−climatic conditions and this may be the cause for variation. The SC is an important trait that is closely associated with the testicular growth and sperm production capacity of domestic animals. Thus, selecting males based on their SC would result in larger testes, potentially with the capacity to produce more spermatozoa (Rounsaville and Foote, 1976; Daudu, 1984).

The stepwise discriminate analysis procedure identified five variables for buck and these are rump height (RH), chest depth (CD), forcanon height (FH), height at wither (HW) and Scrotal length (SL) and six for doe rump height (RH), chest depth (CD), forcanone height (FH), body weight (BW), neck length (NL) and ear length (EL) as most significant discriminating traits. The result was compared with study of Gatew (2015) who explain seven (HL, BW, EL, CG, HW, CW and PW) for does and five (HW, HL PW, CG and EL) for bucks discriminating traits of Bati, Borena and Short Eared Somali Goat Populations.

The relative importance of the identified traits in discriminating both goat populations was assessed at 5% level of significance. Wilk’s Lambda value, the partial R^2^ dropped down as significant discriminating variables added chronologically, describing the amount of variability in each variable accounted by the population differences. As represented by the respective partial R^2^ and F-values; RH was found to have the highest discriminating power in buck followed by CD, FH, HW, and SL in descending order. In the mean time, RH had the highest discriminating power in female followed by CD, FH, BW,NL and EL from the highest to lowest. This implies that bucks required slightly fewer traits measurements to differentiate bucks of the two districts than does which require more variables. This result is inconsistance with report of Gatew (2015) who report HL and HW highest discriminating power in does and buck respectively in Bati, Borena and Short Eared Somali goat Populations.

Body Weight has been the pivot on which animal production thrives. Regression of body weight over quantitative traits, which have higher correlation with body weight, was done to set adequate model for the prediction of body weight separately for each sex. Regression analysis is commonly used in animal research to describe quantitative relationships between a response variable and one or more explanatory variables such as body weight and linear body measurements especially when there is no access to weighing equipment (Cankaya, 2008). To predict the best fitted variables to estimate live body weight and their contribution. Best fitted equation was selected using higher value of adjusted coefficient determination (R^2 adjusted^) which represent the total variability explain by the model and smallar value of mallows C(P) statistics, Akaike information criterion(AIC), root mean square error (RMSE) and Schwarz Bayesian information criterion (SBC) at different age class and sex categories.

In female sampled goat population four variables were positively contributing to the prediction of model which include heart girth, hight at wither, body length and pelvic width were fitted as first, second, third and fourth which account 84% of total variability and heart girth alone also acounts 51% of variation in body weight. Multi linear regression model showed that female had higer adjusted R2 (84%) than male goat population (63%). This indicates that those linear body measurment might predict more accurate in female than male (Bosenu *et al*, 2014). This study shows that heart girth was more reliable in predicting body weight than other linear body measurenments. In this regard, study Thiruvenkadan (2005) described that the better association of body weight with heart girth was possibly due to relatively larger contribution of heart girth, which consists of bones, muscles and viscera.

## Conclusion and recommendation

A systematic description/characterization of the goat types and management systems should be considered as prerequisite for planning the rational use of indigenous goat resources. A coat color pattern varies from population to population depending up on the agro-climatic differences, preferences by their herders and other factors such as the genetic makeup of populations. This study indicates that goats in study area has share common coat color patterns with Woyto Guji and Arsi-Bale goats probably as a result of gene flow between these two neighboring populations. The result in this study also revealed that the smaller mean values for most morphometric measurements dictated the least differentiation between Abaya and Yirgachafee goats. However, a diversity of qualitative traits like coat color, facial and back profile, presence or absence of horn, wattle, ruff and beard was observed among the two goat types. Since the breeders (producers) can easily distinguish desirable phenotypic characteristics, the variability of those traits could be useful in selection program. Due to high and positive correlation coefficients found between body weight and other linear body measurements (HG, BL, HW, HL and PW), selection of one or more of these traits may increase live body weight of these goat populations. Stepwise discriminate analysis procedure was identified RH is highest discriminating power in both does and buck.

The present phenotypic characterization of goats in the study areas has to be further supported with molecular characterization, particularly for their high prolificacy to make use of these peculiar goat populations. Adaptive traits which community acquired through generation have to be improved by applying community based breeding program.

## Authors’ contributions

The author participated in designed all research and wrote the manuscript.

## Availability of data and materials

Data was available in the hands of Corresponding Author

## Ethics approval and consent to participate

The manuscript does not contain clinical studies or patient animals/Goat

## Consent for publication

Not applicable

## References

Belete Asefa. 2013. On farm phenotypic characterization of indigenous goat types and their production system in bale zone of Oromiya region, Ethiopia. M.Sc Thesis Haramaya University,Haramaya,Ethiopia.

Betz, N. E. 1987. Use of discriminate analysis in counseling psychology research. Journal of Counseling Psychology, 34(4): 393

Biruh Tesfahun, Kefelegn Kebede & Kefena Effa. 2017. Traditional goat husbandry practice under pastoral systems in South Omo zone, southern Ethiopia. Tropical Animal Health Production, 49(3): 625–632.

Bosenu Abera, Kefelegn Kebede, Solomon Gizaw and Teka Feyera. 2014. On-Farm Phenotypic Characterization of Indigenous Sheep Types in Selale Area, Central Ethiopia. Veterinary Science and Technology, 5(3):1.

Brown, M. T. and Tinsley, H. E. A. 1983. Discriminate analysis. Journal of Leisure Research, 15(4), 290–310.

Cankaya, S. 2008. A comparative study of some estimation methods for parameters and effects of outliers in simple regression model for research on small ruminants Ondokuz Mayis University, Biometry and Genetics Unit, Kurupelit /Samsun, Turkey.

Daudu, C.S. 1984. Spermatozoa output,testicular sperm reserve and epididymalstorage capacity of the Red Sokoto goats indigenous to northern Nigeria. Theriogenolgy,21(2):317–324.

FAO. 2011. Draft guidelines on phenotypic characterization of animal genetic resources. Commission on genetic resources for food and agriculture, thirteenth regular session,Food and Agriculture Organization of the United Nations, Rome, Italy.

FAO. 2010. Breeding strategies for sustainable management of animal genetic resources. FAO Animal Production and Health Guide lines, Number 3 Rome, Italy.

Farm-Africa. 1996. Goat types of Ethiopia and Eritrea. Physical description and management systems. Published jointly by FARM-Africa, London, UK, and ILRI (International Livestock Research Institute), Nairobi, Kenya.

Hassen, H. 2012. Phenotypic characterization of Ethiopian indigenous goat populations. African Journal of Biotechnology African Journal of Biotechnology Vol. 11(73), pp. 13838–13846

Hirpa, A. & Abebe, G. 2008. Economic Significance of Sheep and Goats Addis Ababa, Ethiopia.

Hussein Hassen. 2015. Phenotypic characterization and breeding practices of Arsi-Bale goat population in selected districts of Arsi and Bale zones, Oromiya regional state, Ethiopia. MSc thesis. Bahir Dar University, Bahir Dar, Ethiopia

Isaac, J.L. 2005. Potential causes and life-history consequences of sexual size dimorphism in mammals. Mammal review, 35(1): 101–115.

Magnabosco, C.D.U., Ojala, M., De los Reyes, A., Sainz, R.D., Fernandes, A. and Famula, T.R. 2002. Estimates of environmental effects and genetic parameters for body measurements and weight in Brahman cattle raised in Mexico. Journal of Animal Breeding and genetics, 119(4):221–228.

Mahilet Dawit. 2012. Characterization of hararghe highland goat and their production system in eastern hararghe. MSc. Thesis Haramaya University, Haramaya, Ethiopia.

Musa, L.M.A., Peters, KJ. and Ahmed, M.K.A. 2006. On farm characterization of Butana and Kenana cattle breed production in Sudan. Livestock research for Rural Development. 18 (12):56–61.

Rege, J.E.O. 1994. Indigenous African small ruminants: a case for characterization and improvement. In: Small Ruminant Research and Development in Africa. Proceedings of the Second Biennial Conference of the African Small Ruminant Research Network. 7-11 December 1992; AICC, Arusha, Tanzania, Addis Ababa Ethiopia: International Livestock Centre for Africa (ILCA)/Technical Centre for Agricultural and Rural Cooperation (CTA).

Rounsaville, T.R. and Foote, R.H. 1976. Heritability of testicular size and consistcy in Holstein bulls. Journal of animal science, 43(1): 9–12.

Semakula, J., Mutetikka, D., Kugonza, R.D. and Mpairewe, D. 2010. Variability in Body Morpho metric measurements and their Application in predicting Live Body Weight of Mubende and Small East African Goat Breeds in Uganda. Middle-East Journal of Scientific Research, 5(2): 98–105.

Tucho, T. A. & Tesfaye, A. 2004. Genetic characterization of indigenous goat populations of Ethiopia using microsatellite DNA markers. NDRI A Doctoral Thesis submitted to the National Dairy Research Institute (Deemed University) Karnal (Haryana), India.

Thiruvenkadan, A.K. 2005. Determination of best-fitted regression model for estimation of body weight in Kanni Adu kids under farmer’s management system. Livestock Research for Rural Development, 17(7):1–11.

Yoseph Mekasha. 2007. Reproductive trait in Ethiopia male goats, with special reference to breed and nutrition. PhD dissertation. Department of clinical sciences, faculty of veterinary medicine and animal science, Swedish university of Agricultural sciences, Uppsala, Sweden.

CSA 2018. Federal Democratic Republic Of Ethiopia Central Statistical Agency report on Livestock and Livestock Characteristics Volume II, Statistical Bulletin 587, Addis Ababa.

FAO 2012. Phenotypic characterization of animal genetic resources. FAO Animal Production and Health Guidelines No. 11. Rome.

FAO 2015 The Second Report on the State of the World’s Animal Genetic Resources for Food and Agriculture, edited by B.D. Scherf & D. Pilling. FAO Commission on Genetic Resources for Food and Agriculture Assessments. Rome (available at http://www.fao.org/3/a-i4787e/index.html).

Gatew, H. 2015. Characterization of indigenous goat populations in selected areas of Ethiopia. American-Eurasian Journal of Scientific Research, 10, 287–298.

Hassen, H. 2012. Phenotypic characterization of Ethiopian indigenous goat populations. African Journal of Biotechnology, 11.

Hirpa, A. & Abebe, G. 2008. Economic Significance of Sheep and Goats.

Takele, A. 2016. Phenotypic characterization of indigenous goat types and their production system in shabelle zone, south eastern Ethiopia. International Journal of Innovative Research and Development, 5, 234–252.

Tesfahun, B., Kebede, K. & Effa, K. 2017. Traditional goat husbandry practice under pastoral systems in South Omo zone, southern Ethiopia. Tropical animal health and production, 49, 625–632.

Tucho, T. A. & Tesfaye, A. 2004. etic characterization of indigenous goat populations of Ethiopia using microsatellite DNA markers. NDRI.

Zergaw, N. & Dessie, T. 2014. On Farm Phenotypic Characterization and Performance Evaluation of Central Highland and Woyto Guji Goat Types for Designing Community Based Breeding Strategies in Ethiopia. Haramaya University.

